# Selective effects of estradiol on human corneal endothelial cells

**DOI:** 10.1101/2023.04.27.538629

**Authors:** Seoyoung Han, Christian Mueller, Caitlin Wuebbolt, Sean Kilcullen, Varinda Nayyar, Brayan Calle Gonzalez, Ali Mahdavi Fard, Jamie C. Floss, Michael J. Morales, Sangita P. Patel

## Abstract

Fuchs endothelial corneal dystrophy (FECD) results from genetic and environmental factors triggering mitochondrial and oxidative stress in corneal endothelial cells (CEnCs) leading to CEnC death and corneal opacification. FECD is more common in women than men, but the basis for this observation is unknown. Because FECD is commonly diagnosed around the time of the menopausal transition in women when estrogen levels decrease precipitously, we studied the effects of the potent estrogen,17-β estradiol (E2) on growth, oxidative stress, and metabolism in primary cultures of human CEnCs (HCEnCs) under conditions of physiologic 2.5% O_2_ ([O_2_]_2.5_) and under hyperoxic stress ([O_2_]_A_: room air + 5% CO_2_). We hypothesized that E2 would counter the stresses of the hyperoxic environment in HCEnCs. HCEnCs were treated ± 10 nM E2 for 7-10 days at [O_2_]_2.5_ and [O_2_]_A_ followed by measurements of cell density, viability, reactive oxygen species (ROS), mitochondrial morphology, oxidative DNA damage, ATP levels, mitochondrial respiration (O_2_ consumption rate [OCR]), and glycolysis (extracellular acidification rate [ECAR]). There were no significant changes in HCEnC density, viability, ROS levels, oxidative DNA damage, OCR, and ECAR in response to E2 under either O_2_ condition. We found that E2 disrupted mitochondrial morphology in HCEnCs from female donors but not male donors at the [O_2_]_A_ condition. ATP levels were significantly higher at [O_2_]_2.5_ compared to [O_2_]_A_ in HCEnCs from female donors only, but were not affected by E2. Our findings demonstrate the overall resilience of primary HCEnCs against hyperoxic stress. The selective detrimental effects of hyperoxia and estradiol on HCEnCs from female but not male donors suggests mechanisms of toxicity based upon cell-sex in addition to hormonal environment.

## Introduction

Fuchs endothelial corneal dystrophy (FECD), the leading indication for corneal transplantation in the United States, is twice as common in women than men, but the underlying pathophysiologic mechanisms for this disparity are unknown [1-4]. Both genetic and extrinsic factors contribute to the development of the common adult-onset form of FECD. A trinucleotide repeat expansion in the *TCF4* gene is commonly associated with FECD in those of European ancestry; however, not all individuals with this expansion demonstrate the disease phenotype and genetics do not explain why FECD is more common in women [5, 6]. Extrinsic stressors thus play an important role in development of FECD.

In FECD, the non-proliferative corneal endothelial cells on the posterior cornea experience oxidative stress, mitochondrial stress, and metabolic stress potentiated by elevated oxygen levels in the anterior chamber of the eye as well as ultraviolet light exposure [7, 8]. These extrinsic stressors contribute to basement membrane abnormalities (guttae in Descemet’s membrane) and corneal endothelial cell death. In mice, ultraviolet-A light exposure is sufficient to generate the FECD phenotype with guttae and corneal endothelial cell loss and greater disease burden in female compared to male mice [9].

We want to understand why FECD is more common in women than men. We focus our study on 17-β estradiol **(E2)**, the most potent circulating estrogen in humans. E2 levels in pre-menopausal women are higher than in men; however, post-menopausal women have lower levels than men [10]. The changes around menopause have been implicated in cardiovascular disease [11], ischemic stroke [12], and ocular surface disease [11-13]. Prior epidemiologic investigations suggest that the peri/post-menopausal age range (∼50-59 years), a common age bracket for diagnosis of FECD, has a disproportionate burden of FECD in women than men compared to other age brackets, suggesting that factors associated with the menopausal transition could be implicated in FECD [14, 15]. Prior laboratory research suggests that estrogen metabolites may be harmful to corneal endothelial cells by forming genotoxic estrogen-DNA adducts and causing DNA damage and apoptotic cell death [16]. However, in studies of cellular and molecular mechanisms of neuronal and cardiovascular diseases, estrogen has protective effects [17-21].

The purpose of our study was to examine the effects of E2 on human corneal endothelial cells **(HCEnCs)** to explore *in vitro* if E2 could be a contributing factor behind the greater prevalence of FECD in women than men. We use a chronic hyperoxic stress model to test the effects of E2 on growth, oxidative stress, and metabolism in primary HCEnC cultures.

## Methods

### Corneal tissues

The use of human tissues for these studies was approved by the University at Buffalo and VA Western NY Healthcare System Institutional Review Boards and research protocols were approved by the VA Western NY Research and Development Committee. Human cadaver corneal tissues were obtained from the Anatomical Gift Program at the University at Buffalo. Human surgical FECD samples from endothelial keratoplasty were obtained from patients following written informed consent. Age and sex of all tissues used for these experiments are listed in Table 1. All experiments, including patient enrollment, were performed between 2016-2023.

**Table 1.**
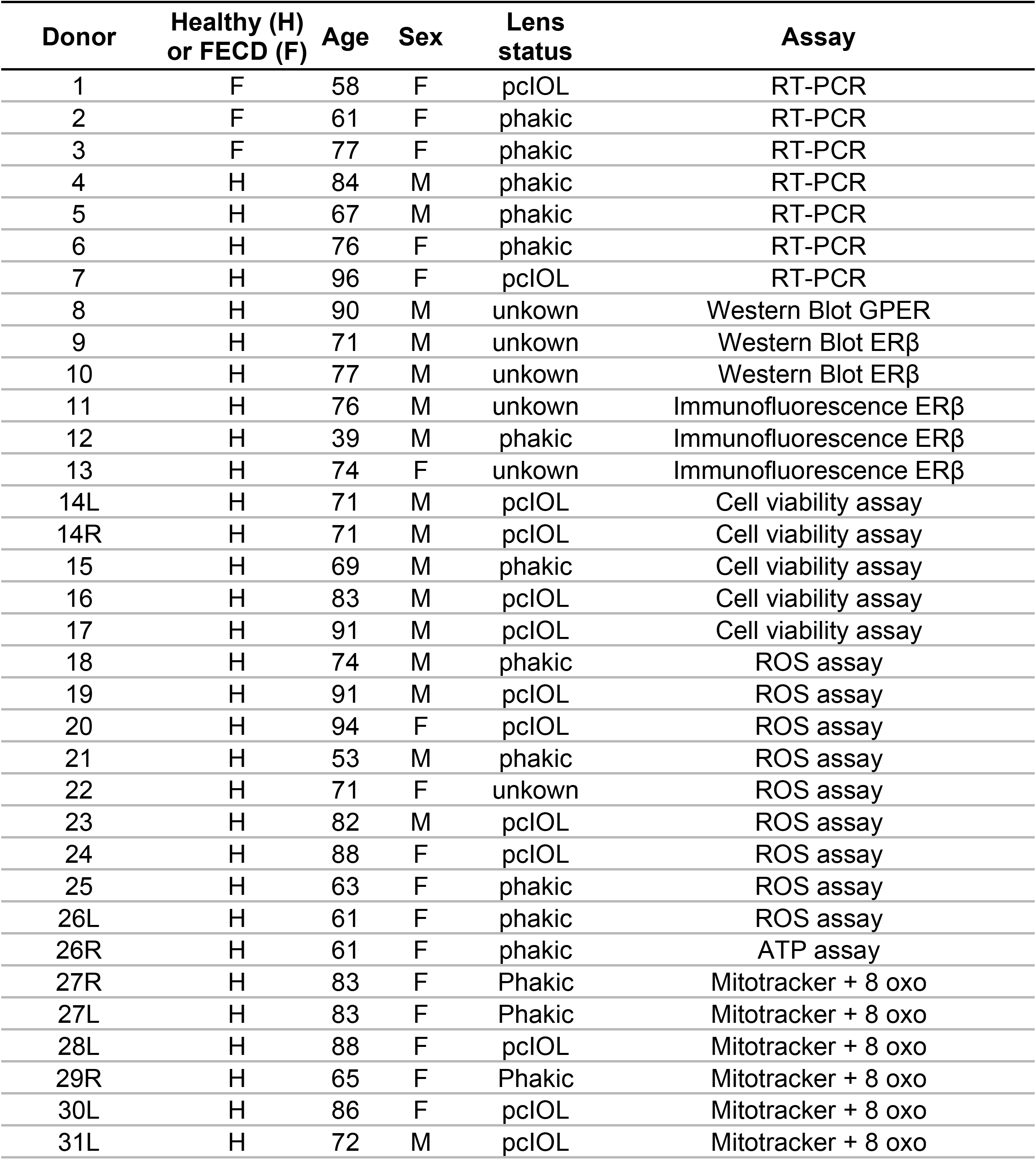
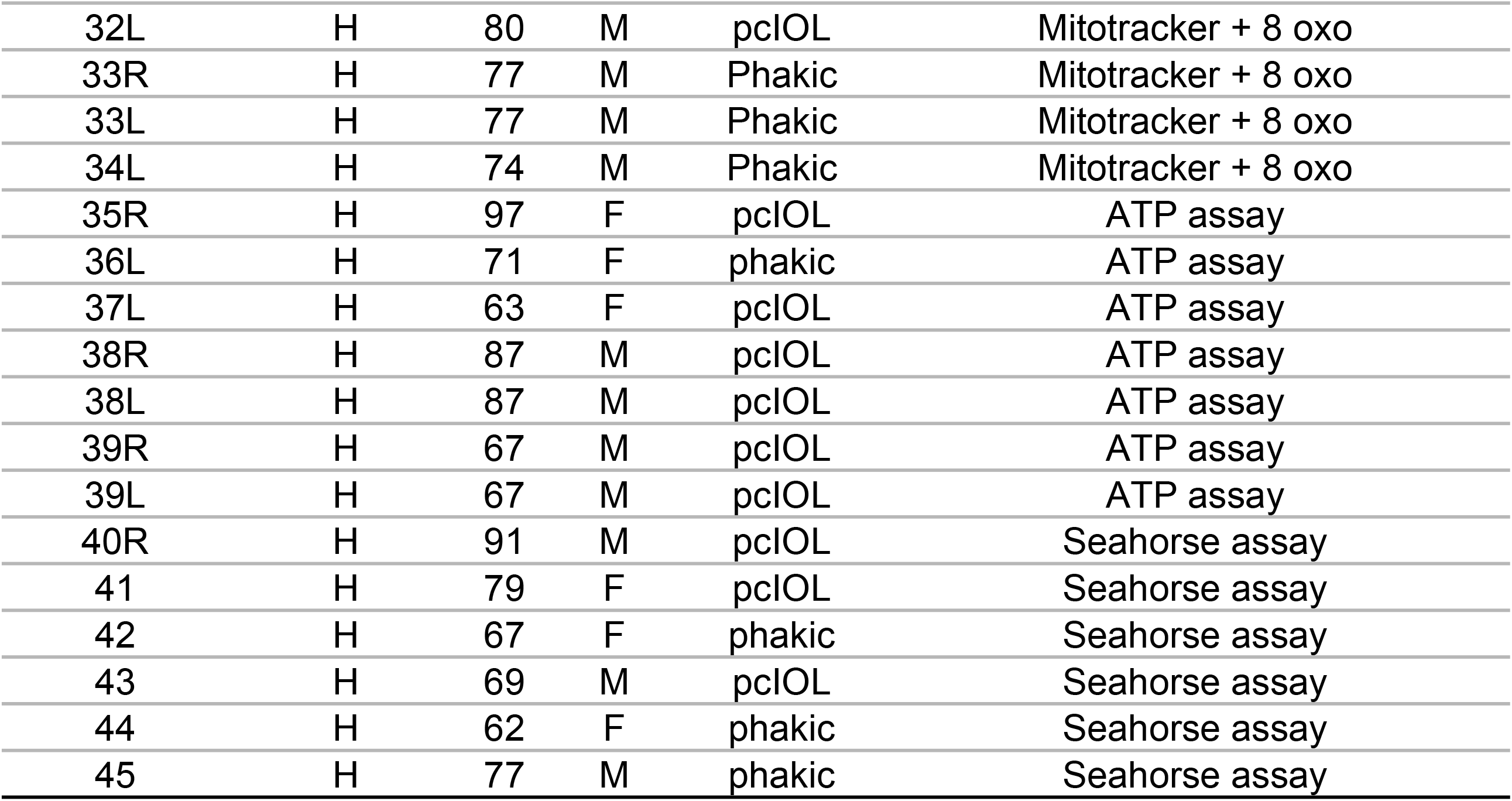
Corneal tissue donor information.

### Cell cultures

All studies of HCEnCs were performed with primary, P0, corneal endothelial cell cultures. Cultures were established using previously published protocols [22]. P0 corneal endothelial cell cultures were initiated from human corneas obtained within 24 hours post death. Corneoscleral buttons were dissected from the eyes and were examined for the presence of guttae. Corneas with guttae were excluded in the experiments. Descemet’s membrane with adherent endothelial cells were stripped from the corneoscleral buttons and incubated overnight at 37°C in minimal medium (Human Endothelial SFM, Gibco, cat# 11111-044); 2% charcoal stripped fetal bovine serum (FBS); 1x antibiotic/antimycotic (Gibco, cat# 15240-062). Cells were dissociated with 0.02% EDTA (Sigma, cat# 45-E8008). Cells were resuspended in growth medium (Opti-MeM1, Gibco; 8% FBS; 20 µg/mL ascorbic acid; calcium chloride 200 mg/L; 1 x antibiotic/antimycotic; 100 µg/mL pituitary extract; 5 ng/mL epidermal growth factor; 50 µg/mL gentamicin) and distributed on FNC-coated (Athena, cat# 0407) plates or coverslips according to the experimental plan.

Cultures were incubated at 37°C in standard room air incubators ([O_2_]_A_: room air + 5% CO_2_ humidified incubator), or at 2.5% O_2_ in a modular incubation chamber ([O_2_]_2.5_, 2.5% O_2_ + 5% CO_2_ + balance N_2_, humidified; Billups-Rothenberg, Inc., Del Mar, CA, USA). HCEnCs were expanded in growth medium to confluence. Following confluence, cells were matured to a stable phenotype in minimal medium. Charcoal stripped FBS was used in the minimal medium to remove endogenous steroid hormones in serum. For experiments testing estradiol exposure, 17β-estradiol (**E2**; stock concentrations selected for preparation in dimethylsulfoxide with 500-fold dilution into culture medium to achieve final desired concentrations; Sigma, cat# E8875) was added for 7-10 days in the maturation phase of the HCEnC culture.

MCF7 and PC3 cells were shared by colleagues who purchased from ATCC (American Type Culture Collection, Manassas, VA, USA) [23, 24]. MCF7 cells were cultured in DMEM (Corning, cat# 10-017-CV) + 10% FBS + 1x antibiotic-antimycotic. PC3 cells were grown in RPMI 1640 medium (Corning, cat# 10-040-CV) with 10% FBS and 1 × antibiotic/antimycotic. Cells were maintained in T25 flasks in a 5% CO_2_, humidified 37°C tissue culture incubator.

### Quantitative RT-PCR

Transcript levels of estrogen receptor (ER) α, β, and G protein-coupled estrogen receptor (GPER) 1 were measured by reverse transcription-real time quantitative polymerase chain reaction (RT-qPCR). RNA was purified from individual corneas using RNeasy Plus Universal kit (Qiagen, Hilden, Germany). cDNA was synthesized using the iScript Advanced cDNA synthesis kit for RT-qPCR (Bio-Rad Laboratories, Hercules, CA, USA) with 2.0-20 ng input corneal RNA according to a manufacturer protocol. Real time qPCR was performed using the Bio-Rad SsoAdvanced Univ SYBR Green kit with 2 μL cDNA (10% total volume) in a 96-well plate. Reactions were performed on a Bio-Rad C1000 thermal cycler equipped with CFX96 Real-Time System running CFX Manager software (version 3.1). Assays were run in triplicate and included controls for genomic DNA contamination, PCR efficiency, and reverse transcription. Primers were designed and validated by the manufacturer (PrimePCR™ SYBR® Green Assay, Bio-Rad) and are listed in S1 Table. The quantification cycles (Cq) were calculated by CFX Manager.

### Western blots

Western blots were performed similarly to published protocols [22]. Protein was isolated from confluent PC3 and MCF7 cultures. For human cornea samples, Descemet’s membrane-endothelium complex was stripped and stored at -80°C and thawed in lysis buffer. Samples were disrupted mechanically with a pestle in lysis buffer (50 mM Tris (pH 7.4), 250 mM NaCl, 2 mM ethylenediaminetetraacetic acid (pH 8.0), 10% Protease Inhibitor Cocktail (Sigma-Aldrich), and 10% Triton X-100). Protein concentration was measured with the Pierce BCA kit (Thermo Fisher Scientific, Waltham, MA, USA) and 10 μg protein was loaded per lane on Mini-PROTEAN TGX Precast Gels (Bio-Rad Laboratories, Hercules, CA, USA). Following protein separation, samples were transferred to PVDF membranes (Bio-Rad tank blotting system), blocked with 10% goat serum and then incubated with primary antibodies (mouse anti-ERβ antibody, #SC-390243, lot# K1814, Santa Cruz Biotech, used at 1:250 dilution in blocking buffer; rabbit anti-GPER1 antibody, cat # HPA027052, lot# D118286, Sigma Aldrich, used at 1:500 diluted in blocking buffer) overnight at 4°C. Following three washes in TBST, secondary antibodies were applied (goat anti-rabbit IgG-Alkaline Phosphatase, A9919, Sigma Aldrich; goat anti-mouse IgG-Alkaline Phosphatase, A4312, Sigma Aldrich; 1:3000) for 1 hour at room temperature. Signals were developed with ECF substrate (Cytiva Amersham, Marlborough, MA, USA) and imaged with the ChemiDoc MP Imaging System (Bio-Rad).

### Estrogen receptor immunofluorescence localization

The Descemet’s membrane-endothelium complex was stripped from fresh human corneas and fixed immediately in 3% paraformaldehyde at 4°C. The tissue was rinsed three times with phosphate-buffered saline solution (PBS) and then permeabilized with 0.1% Triton X-100 for five minutes and washed two times with PBS. Following block of non-specific antibody binding with 10% goat serum for 30 minutes at room temperature, mouse anti-ERβ (1:100) was added and incubated for 2 hours at room temperature. Antibody solution was then removed, the sample was washed with PBS three times for five minutes, and incubated at room temperature for 1 hour with goat anti-mouse IgG conjugated to Alexa Fluor 568 (1:250, Invitrogen, cat # A11011). The antibody solution was removed, and the sample was washed three times with PBS for five minutes. The tissue was placed carefully in a single layer onto a microscope slide containing a drop of Vectashield with DAPI (Vector Laboratories, Burlingame, CA, USA). Samples were imaged by laser confocal microscopy (Leica TCS SPEII DMI4000).

### Estrogen receptor agonist experiments in PC3 cells and HCEnC

PC3 cells between passage 12-15 were used for experiments. PC3 cells at 70% confluency were collected and seeded at 30,000 cells per well in a 24-well plate with culture medium containing charcoal stripped fetal bovine serum. After 24 hours, each well was treated with 1, 10, or 100 nM E2. Similarly, other wells were treated with 0.1, 1.0, or 10 µM G1 GPER agonist (Azano Biotech, Albuquerque, NM) dissolved in culture media. Cells were incubated for 3 days without a change of culture medium. Cells were then lifted with trypsinization, stained with trypan blue, and viable cells from each well were counted by hemocytometer. Data from 4 experiments were averaged and compared with t-tests assuming equal variance.

For HCEnCs, cells from one cornea were distributed into 16 wells (8 at [O_2_]_A_ and 8 at [O_2_]_2.5_ in 96-well plates). Once cells were confluent, they were switched to minimal medium (with charcoal stripped fetal bovine serum) with or without E2 or G1 (2 wells at each condition). Following incubation for 7-10 days, cells were then stained with DAPI and nuclei were counted to determine the total cell count. The mean data were compared by ANOVA with significance at *p* < 0.05.

### Cell viability assays

HCEnC cultures were cultured in two 96-well plates, with cells from one cornea distributed into 16 wells, 8 wells at [O_2_]_A_ and 8 wells at [O_2_]_2.5_. The cells were grown in growth medium until confluence, and then matured in minimal medium. During the maturation phase, cells were grown in different conditions, 2 wells for each condition: control, 1 μM GPER agonist (AZ00004-G1, Azano Biotech, Albuquerque, NM), 10 nM E2, 10 nM E2 + 100 nM GPER antagonist (AZ00001-G36, Azano Biotech, Albuquerque, NM). Two wells with minimal medium without cells were used for negative control. Cell viability was measured using the RealTime-Glo Cell Viability Assay (Promega, Madison, WI, USA). MT Viability Substrate and NanoLuc Enzyme at 1:2000 dilution were added to each well, and luminescent signal was measured with a Synergy HT plate reader (BioTek, Winooski, VT). After the reading, cells were stained with DAPI and number of nuclei counted. Luminescence readings were normalized to cell count per well. The mean data were compared by ANOVA with significance at *p* < 0.05.

### Measurements of cellular reactive oxygen species

Primary HCEnCs from one cornea were distributed into four wells (two wells each in two 24-well plates). Cultures were maintained under [O_2_]_A_ and [O_2_]_2.5_. Cells were fed with growth medium until confluence and then matured for 7-10 days in minimal medium with or without 10 nM E2 for the duration of maturation. Supernatant from each plate was removed and HCEnCs were immediately lysed with 0.5% Triton X-100 in PBS. Lysed cells were then transferred to microfuge tubes, incubated on ice for twenty minutes, centrifuged at 4°C for 20 minutes to pellet cell debris, and the supernatant was collected for analysis. ROS levels were analyzed with the In Vitro ROS/RNS Assay (Cell Biolabs San Diego, CA). 35 μL of cell lysate mixed with 15 μL PBS were added to wells of a 96-well plate. 100 μL of dichlorodihydrofluorescin solution, prepared according to manufacturer instructions, were added to each well and incubated for thirty minutes protected from light. Fluorescence was measured using a fluorescence plate reader (SynergyTM HT, BioTek Instruments, Inc, Winooski, VT) at 480 nm excitation and 530 nm emission for five minutes. Fluorescence measurements from cell lysates were compared to a hydrogen peroxide standard curve and normalized to total protein concentration in each sample. The protein concentration of the lysate was determined with the Pierce BCA Kit. The mean data were compared by ANOVA with significance at *p* < 0.05.

### Mitotracker and 8-hydroxy-2’-deoxyguanosine (8-oxo-dG) immunofluorescent staining

HCEnCs were seeded in 48-well plates (one cornea per 8 wells) with round #1.5 glass coverslips. Cells were grown to confluence in growth medium and then switched to minimal medium with or without 10 nM E2 for 1-2 weeks prior to 8-oxo-dG assay. 48 and 24 hours prior to the staining protocol, 100 μM H_2_O_2_ was added directly to culture medium in the desired wells. At the time of assay, HCEnCs were incubated in 200 nM MitoTracker Red CMXRos (Invitrogen, M5712) by adding 1.0 μl of 50 μM stock in each well containing 250 µL of minimal medium for 15 minutes at 37°C. Cells were then fixed with 1:1 methanol:acetone for 20 minutes at -20°C. Methanol:acetone was then removed, and samples were allowed to air dry, followed by sequential washes with PBS, 35%, 50% and 75% ethanol in H_2_O for 3 minutes each. Next, DNA was denatured by incubation of cells with 0.15 N sodium hydroxide prepared in 70% ethanol for 4 minutes, followed by 3.7% paraformaldehyde in 70% ethanol for 2 minutes. Samples were then washed with 50% ethanol, 35% ethanol, and PBS for 2 minutes each. Cells were permeabilized in 0.1% triton-X100 for 5 minutes and washed with PBS twice for 5 minutes. Non-specific binding was blocked with 10% normal goat serum in PBS for 30 minutes at room temperature, and then incubated with anti-8-oxo-dG antibody (1:400 in 1% goat serum; R&D Systems, 4354-MC-050, Lot# P275856) for 1 hour. Following 3 x 5-minute washes in PBS, cells were incubated in secondary antibody (goat anti-mouse IgG conjugated to Alexa Fluor 488) diluted 1:1000 in PBS containing 1% goat serum for 1 hour at room temperature. Cells were again washed with PBS for 3 x 5 minutes with a final wash with de-ionized water. The samples were then mounted with Vectashield with DAPI. Samples were imaged by fluorescence microscopy with a Keyence scanner (Keyence BZ-X810; Keyence, Osaka, Japan) with identical exposure and acquisition settings. From the digital images, a minimum of 50 cells for each sample well were analyzed. The mitochondrial morphology for each cell was graded as diffuse (normal), intermediate, or fragmented (abnormal) [25]. 8-Oxo-dG was graded by nuclear stain as positive or negative. Mitochondrial morphology and 8-oxo-dG were graded by masked observers. Mean data were analyzed by ANOVA with significance at *p* < 0.05. Significant findings were explored with Tukey post-hoc test.

### ATP assay

HCEnCs from each cornea were divided into in two 24-well plates, two wells each (one cornea per 4 wells) and were grown to confluence under [O_2_]_A_ and [O_2_]_2.5_ in growth medium and were then treated in minimal medium with or without 10nM E2 for 7-10 days. ATP concentration was measured using a luminescent ATP detection kit (Abcam, Cambridge, MA) according to manufacturer instructions. First, 50 μL of kit detergent was added to each well of HCEnCs. The plate was sealed and placed on an orbital shaker at 600-700 rpm for 5 minutes to lyse cells and stabilize ATP. The cell lysate was transferred to a 96-well plate. 50 μL of substrate solution was added to each well, and the 96-well plate was shaken again on the orbital shaker for 5 minutes. The plate was dark adapted for 10 minutes prior to luminescence measurement on a plate reader (Synergy HT). Raw luminescence values were compared to a standard curve run in parallel with the assay in order to calculate ATP concentration for each well. Mean ATP concentrations measured from each condition were compared by ANOVA and Tukey’s test with significance at *p* < 0.05.

### Seahorse assay

Seahorse XFe24 analyzer (Agilent Technologies, Inc., Santa Clara, CA, USA) was used to measure O_2_ consumption rates (OCR; reflective of mitochondrial respiration) and extracellular acidification rates (ECAR; reflective of glycolysis). Primary HCEnCs were dissociated and seeded directly into FNC-coated Seahorse XFe24 culture plates. Cells from one donor cornea were seeded into 10 wells equally distributed in two plates, one plate incubated at [O_2_]_A_, and one plate incubated at [O_2_]_2.5_. Cells were expanded until confluence in growth medium and then matured 7-10 days in minimal medium. For half of the wells, the incubation in minimal medium included addition of 10 nM E2 and the other half served as control without E2. The Seahorse XFe24 sensor cartridges and drugs were prepared according to manufacturer protocols as previously described by our laboratory [22]. The assay medium was Seahorse XF Dulbecco’s modified Eagle medium, with 1.2 mM glutamine, 7.0 mM glucose, and 0.45 mM pyruvate. OCR and ECAR measurements were performed at baseline and after additions of oligomycin (1 μM), FCCP (1.5 μM), rotenone/antimycin A (0.5 μM), and 2-DG (50 mM; all drugs were purchased from Sigma-Aldrich). Following completion of the assay, cells were fixed with 1:1 methanol:acetone at −20°C, stained with DAPI, imaged, and cell nuclei were counted to normalize OCR and ECAR to cell density. Trends in mean data values following each Seahorse assay drug addition were compared between control and E2 groups at both [O_2_]_A_ and [O_2_]_2.5_ using two-tailed t-tests.

## Results

### Estrogen receptors in human corneal endothelium

To determine if estrogen could affect the corneal endothelium by canonical estrogen receptor signaling pathways, we investigated the expression of estrogen receptors in native human corneal endothelium from males and females with and without FECD. Using quantitative RT-PCR, we found that the G-protein coupled estrogen receptor (*GPER*) was most abundantly expressed in normal corneal endothelium followed by estrogen receptor β (*ERβ*) (Fig 1). Estrogen receptor α showed very low expression. The same pattern of estrogen receptor expression was noted in corneal endothelium from FECD patients. We further explored GPER and ERβ protein expression. The GPER antibody detected multiple distinct bands on Western blot of protein from human corneal endothelium, PC3 cells, and MCF7 cells (Fig 2A); however, none of the bands corresponded to the theoretical molecular weight of GPER (∼42 kDa) suggesting alterative splicing of GPER transcripts, post-translational modifications of GPER protein, or antibody detection of a similar antigen in different proteins. Due to this variability, further testing was not conducted for GPER. ERβ antibody detected a single protein of expected molecular weight (55-60 kDa) in human corneal endothelium and PC3 cells, but not in MCF7 cells where ERα is much greater than ERβ (Fig 2B). ERβ was also present in native corneal endothelium, with most cells having cytoplasmic expression and some having nuclear expression (Fig 2C). These data demonstrate that estrogen receptors are present in human corneal endothelium and human corneal endothelium could thus be susceptible to receptor-mediated estrogen signaling.

**Fig 1.**
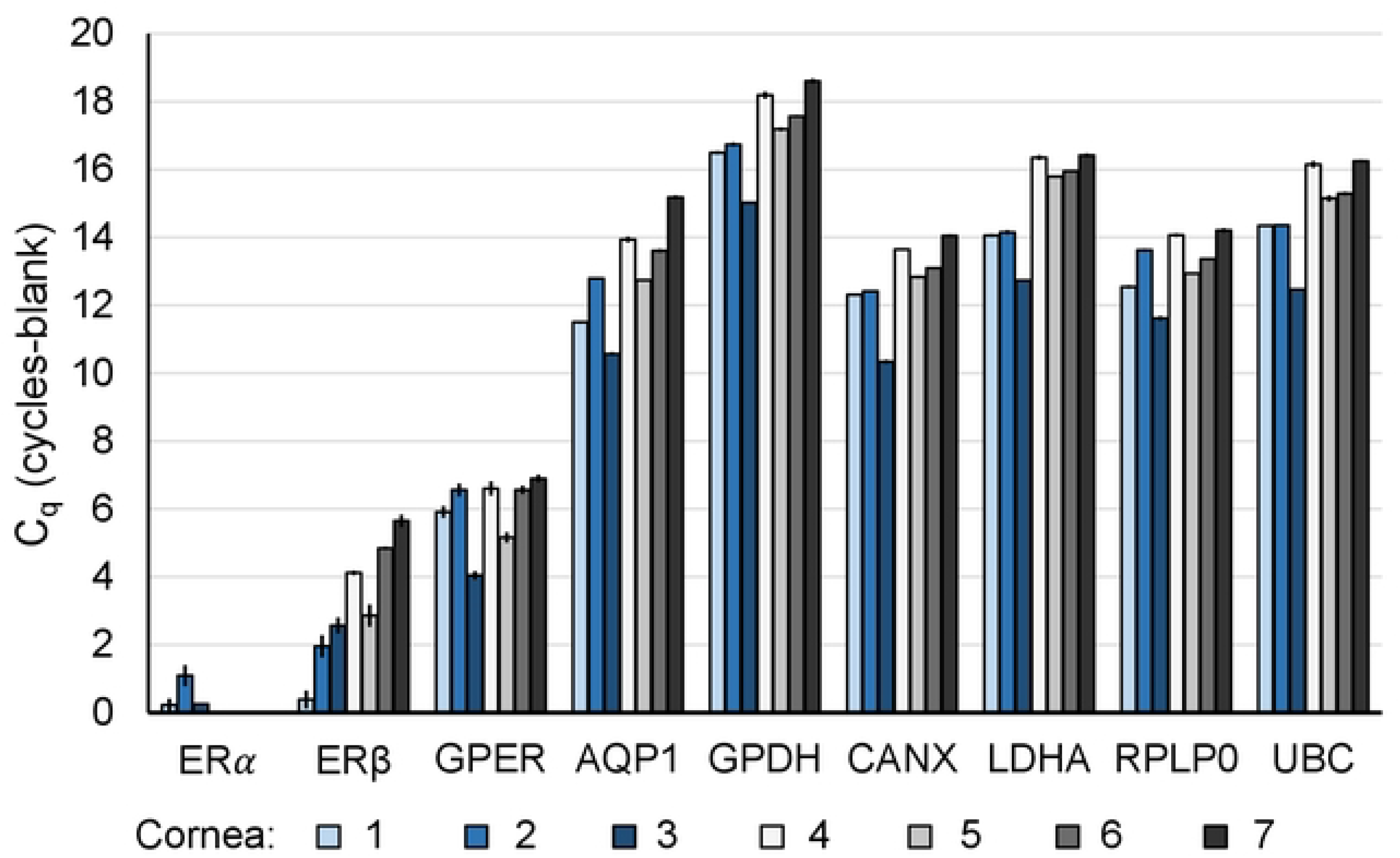
Estrogen receptor gene expression in human corneal endothelium by qRT-PCR. mRNA expression for *ERα*, *ERβ*, and *GPER,* (and housekeeping/control genes *AQP1*, *GPDH*, *CANX*, *LDHA*, *RPLP0, and UBC*) in non-diseased human corneal endothelium (n= 4, corneas #4-7), and FECD patient corneal endothelium (n=3, corneas #1-3). Data presented are the mean of three reverse transcription reactions ± standard deviation. Donor numbers correspond to those in Table 1.

**Fig 2.**
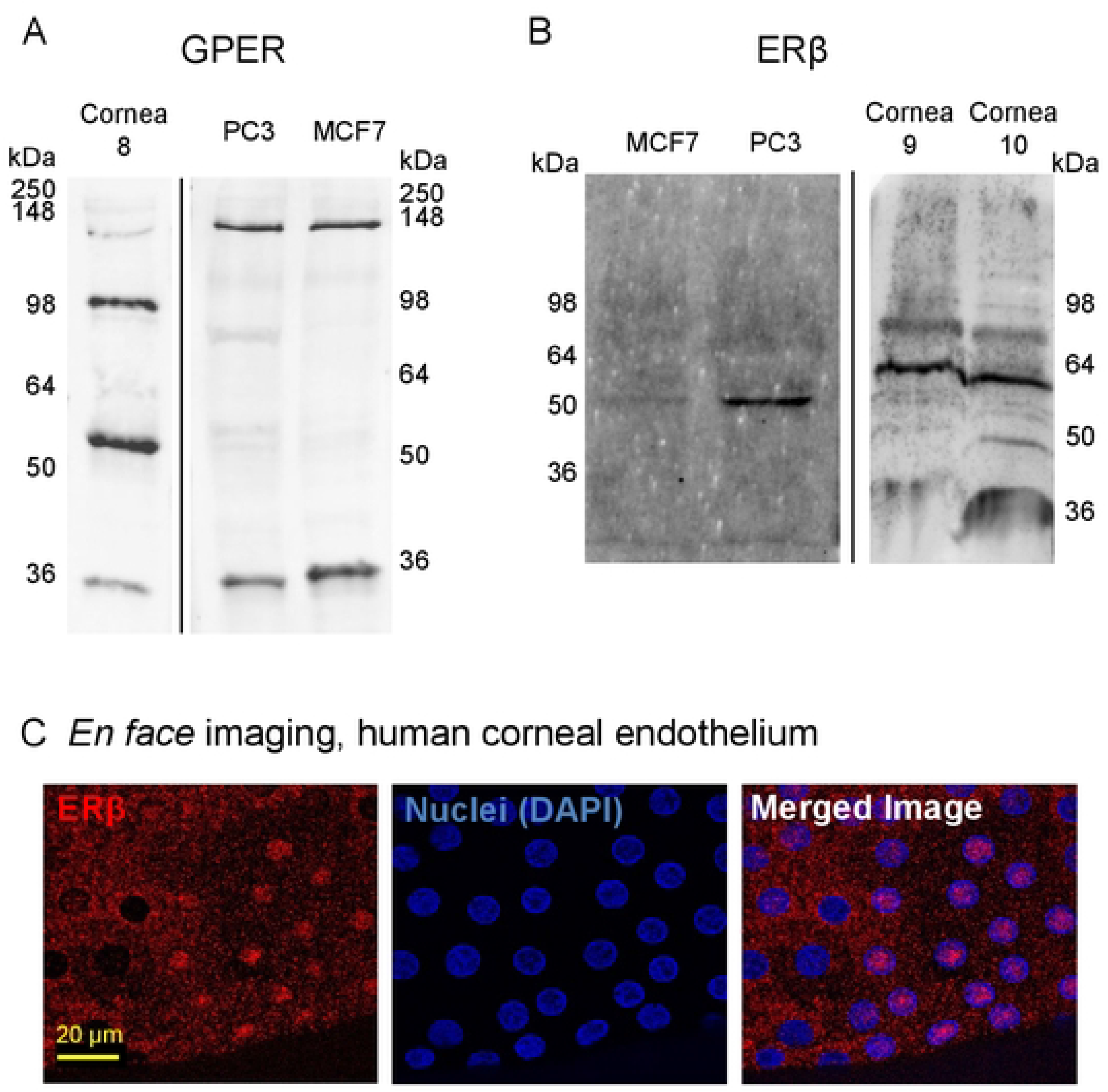
Estrogen receptor protein expression in human corneal endothelium. Western blot for: (A) GPER and (B) ERβ in native human corneal endothelium (n=1 donor for GPER, cornea #8; n=2 donors for ERβ, corneas #9, 10), and PC3 (n=3 for each antibody), and MCF7 (n=3 for each antibody) cells lines. Theoretical molecular weight of GPER is 42 kDa and ERβ is 55-60 kDa. (C) *En face* immunofluorescence localization of ERβ in normal human corneal endothelium (representative images from corneas #11-13). Donor numbers correspond to those in Table 1.

### Effects of E2 and GPER agonist on HCEnC growth

Estrogens can promote cell growth or cell death depending upon the cell type or environment of the cell [26, 27]. We studied the effect of E2 on primary cultures of HCEnCs. In addition, because GPER is the most abundant estrogen receptor transcript in corneal endothelium, we studied the effects of the GPER agonist, G1.

To establish the concentrations of G1 and E2 to use in experiments, we evaluated the effects of both drugs on PC3 cells. Both E2 and G1 treatment of PC3 cultures resulted in a significant decrease in PC3 cell numbers (Fig 3A). For the remainder of experiments described here, we used G1 at 1 μM and E2 at 10 nM.

**Fig 3.**
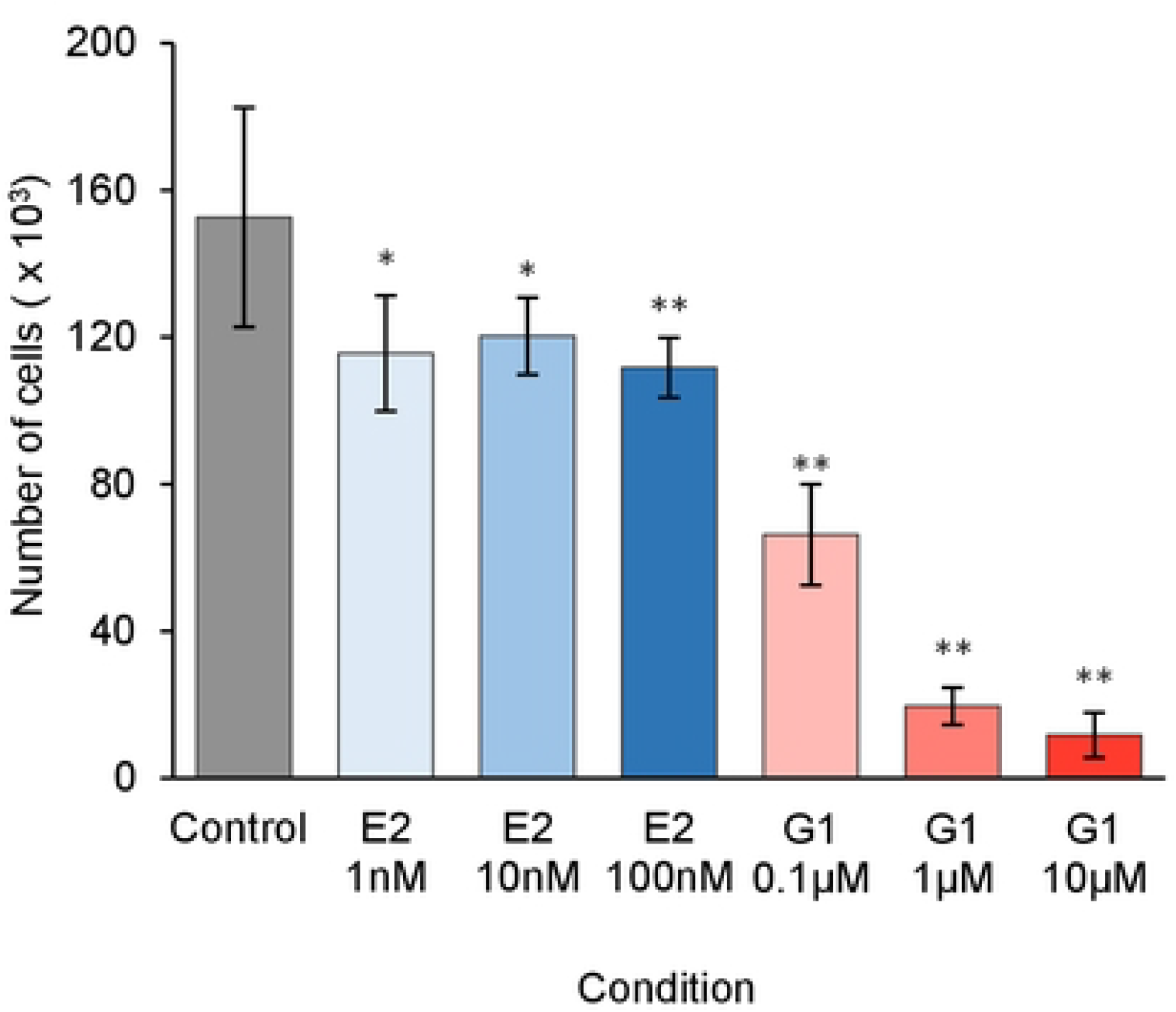
Effects of G1 and E2 on PC3 and HCEnC on cell growth. (A) Number of PC3 cells with 1 nM, 10 nM, and 100 nM E2 or 0.1 µM, 1.0 µM, 10 µM G1 treatment (n=12). *P*-values from t-test compared to the control: ∗ *p*<0.01, ∗∗ *p*<0.001. (B) HCEnC counts with 10 nM E2 or 1.0 μM G1 treatment (n=5, corneas #14-17). Data presented as mean ± SD of total cell counts per well. *P*-values from ANOVA single factor.

We evaluated the effects of E2 and G1 on HCEnC growth. The HCEnCs in these experiments were cultured under physiological O_2_ conditions [28] with 2.5% O_2_ ([O_2_]_2.5_) and under the stress of chronic high oxygen conditions in a room air culture incubator ([O_2_]_A_) [22]. These two conditions were chosen in order to model the oxygen stress seen in FECD [29]. We found that in contrast to the effects on PC3 cells, neither E2 nor G1 affected the growth of HCEnCs cultured under either O_2_ condition (Fig 3B).

### E2 and G1 effects on HCEnC viability

Because E2 can affect more than just the growth of cells, we further investigated the effects of E2 on various functions of HCEnC. In particular, we were interested to see if E2 could affect the viability of HCEnC in stressed environments and if G1 could reproduce the effects. We used the RealTime-Glo MT cell viability assay to measure the reducing potential of HCEnC in the presence and absence of oxygen stress, E2, and G1. This assay was chosen because it was a non-terminal assay that would allow measurements at multiple time points. Baseline measurements were performed following 4 days of incubation with E2 or G1 at either [O_2_]_A_ or [O_2_]_2.5_ (Fig 4A). There were no statistically significant differences compared to control. We next evaluated if E2 or G1 would affect viability upon the stress of reversal of O_2_ culture condition. For this experiment, cultures at [O_2_]_A_ were placed at [O_2_]_2.5_ and cultures at [O_2_]_2.5_ were placed at [O_2_]_A_. The stress of O_2_ condition change had variable effects on cells from different corneas (Fig 4B-G), but there were no statistically significant differences in the response to stress under any condition.

**Fig 4.**
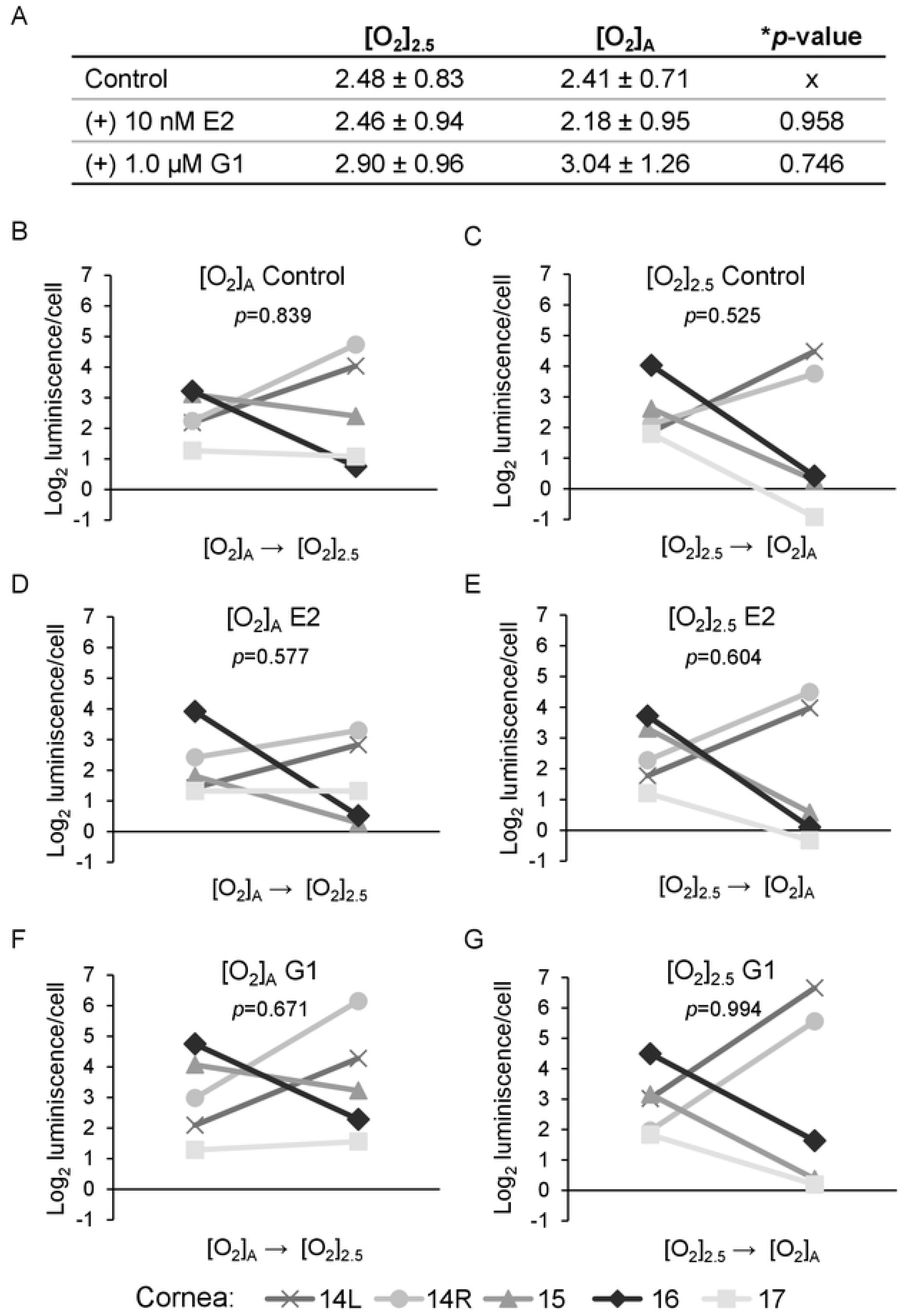
Cell viability assay of HCEnC in the presence and absence of oxygen stress, and E2 and G1 treatment. (A) Cell viability measurements (log_2_ luminescence/cell) with E2 or G1 treatment in HCEnCs (n=5, corneas #14-17). Data is presented as mean ± SD. \**p*-value from ANOVA. Each plot in B-G represent the data for a single donor with *p*-value from paired t-tests. Donor numbers correspond to those in Table 1.

### Oxidative stress

Oxidative stress from oxidant-antioxidant imbalance leading to the accumulation of oxidized DNA lesions is a key finding in FECD [7]. Furthermore, mitochondrial dysfunction and fragmentation from increased reactive oxygen species are hallmarks of FECD that lead to corneal endothelial cell death [30, 31]. In many cell types, oxidative stress is regulated by E2 [32, 33]. We therefore examined the role of E2 in mediating the effects of hyperoxic stress in HCEnCs by measuring reactive oxygen species (ROS), changes to mitochondrial morphology, and oxidative DNA damage.

Levels of total ROS in primary HCEnC cultures were measured with an In Vitro ROS/RNS Assay (Cell Biolabs). ROS levels were unaffected by culture O_2_ conditions and were unchanged in the presence or absence of E2 (Table 2). There were no differences in effects between males and females.

**Table 2.** Levels of ROS in HCEnCs.

|  |  | [O<sub>2</sub>]<sub>2.5</sub> |  | [O<sub>2</sub>]<sub>A</sub> |  | *p-value |
| --- | --- | --- | --- | --- | --- | --- |
|  |  | (-) E2 | (+) E2 | (-) E2 | (+) E2 |  |
| ROS<br>(nM ROS/μg protein) | Males | 55.80 ± 32.99 | 32.62 ± 15.34 | 24.70 ± 15.15 | 43.44 ± 34.43 | 0.42 |
|  | Females | 19.45 ± 6.68 | 28.31 ± 9.72 | 23.94 ± 17.57 | 31.44 ± 8.56 | 0.40 |
|  | Total | 40.22 ± 30.60 | 30.22 ± 11.86 | 24.45 ± 14.13 | 36.77 ± 22.83 | 0.53 |

Data presented as mean ± SD \*p-value from ANOVA n= 9 total, 4 males, and 5 females. Corneas #18-26L Donor numbers correspond to those in Table 1

We next evaluated qualitative changes in mitochondrial morphology. Mitochondria were stained with MitoTracker Red CMXRos and the mitochondrial arbor morphology was classified in each cell as diffuse (normal), fragmented (abnormal), or intermediate (Fig 5) [34]. A positive control with 100 μM H_2_O_2_ showed an expected significant decrease in the percentage of cells with diffuse mitochondrial arbor and increase in cells with fragmented mitochondrial arbor (Fig 6A-C). When the HCEnCs were exposed to [O_2_]_A_ stress in the presence and absence of E2, we found a significant decrease (*p*=0.019, single factor ANOVA) in the percentage of cells with diffuse mitochondrial arbor and increase in cells with intermediate mitochondrial morphology (*p*=0.007; Fig 6D). Post hoc analysis revealed that the significant decrease in cells with diffuse mitochondria was seen only in cells at [O_2_]_A_ with 10 nM E2 compared to [O_2_]_A_ without E2 (*p*=0.039). The post-hoc tests for intermediate morphology for the total experimental population had no physiologically relevant differences. However, when the intermediate morphology data were divided by the sex of the corneal donor tissue, the data were significant for an increase in cells with intermediate mitochondrial morphology in females at [O_2_]_A_ with 10 nM E2 compared to [O_2_]_A_ without E2 (*p*=0.044). There were no significant differences in males. There were no differences in mitochondrial morphology due to the hyperoxic stress alone. These data show that detrimental changes to mitochondrial morphology in HCEnCs occur in response to hyperoxic stress only in the presence of E2 in females.

**Fig 5.**
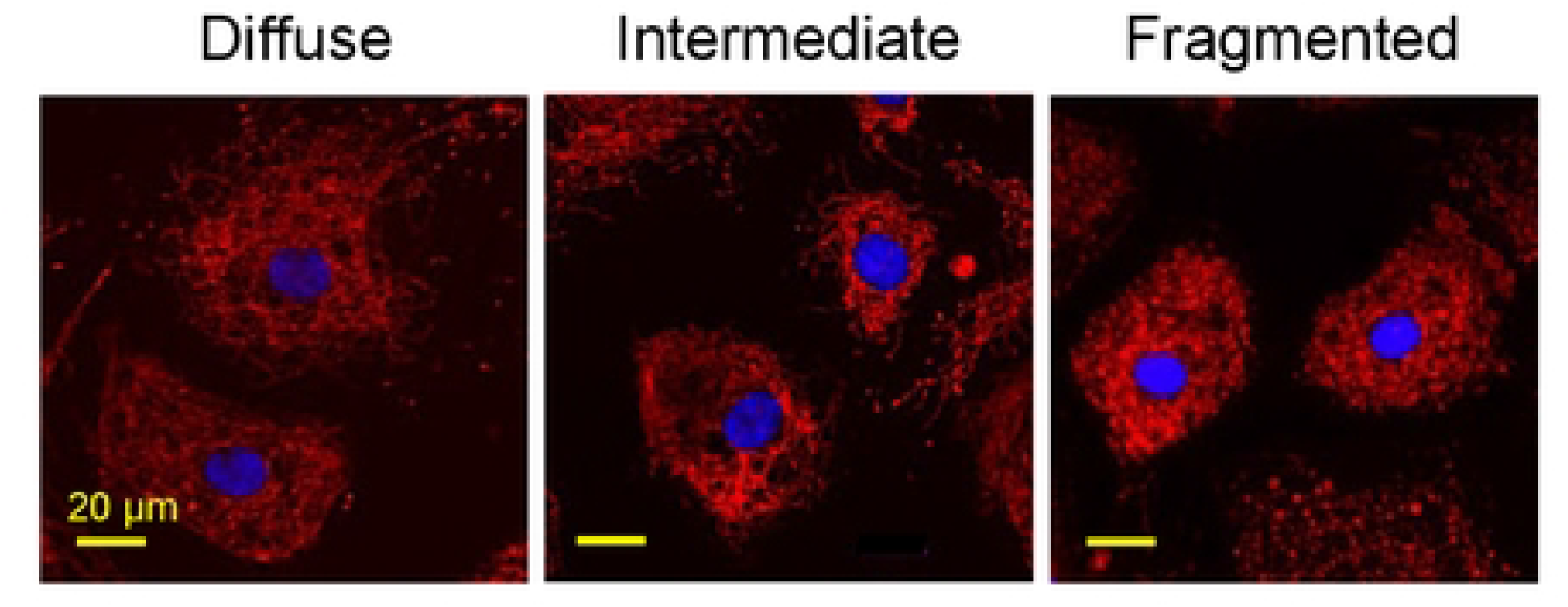
Representative images of mitochondrial morphology grading. Mitochondrial arbor from MitoTracker Red stain was graded for each cell as diffuse (normal), intermediate, or fragmented (abnormal). Nuclei were stained with DAPI (blue). Representative images from corneas #27R-34L.

**Fig 6.**
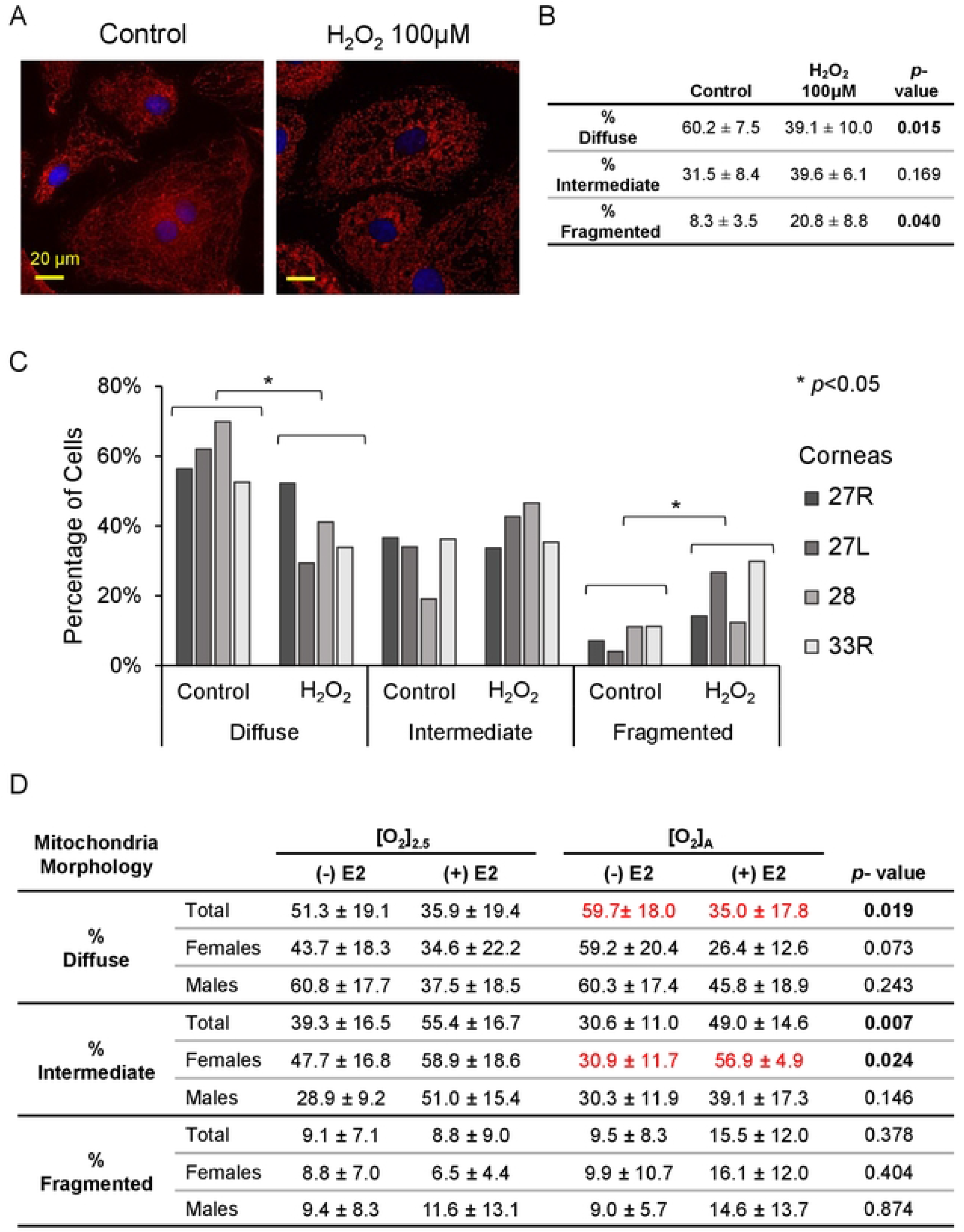
Mitochondrial morphology in HCEnCs in the presence and absence of oxygen stress, and estradiol treatment. (A) Immunofluorescence images of mitochondrial arbor, stained with MitoTracker Red, in HCEnCs following H_2_O_2_ treatment. (B-C) Quantification of mitochondrial arbor grade as diffuse (normal), intermediate, or fragmented (abnormal) following H_2_O_2_ treatment in HCEnCs (n=4, corneas #27R, 27L, 28L, 33R). *P*-value from t-test compared to the control. (B) Data presented as mean ± SD. (C) Donor numbers correspond to those in Table 1. (D) Quantification of mitochondrial morphology grade with the presence or absence of 10 nM estradiol treatment at [O_2_]_2.5_ or [O_2_]_A_. *P*-value from ANOVA single factor. Pairwise comparisons were performed by Tukey post hoc analysis, with significant differences highlighted in red. N= 9 total, 4 males, and 5 females (corneas # 27R-33L).

Mitochondrial stress can also result in cellular oxidative stress with oxidative DNA damage. We tested for the oxidative DNA damage marker, 8-hydroxy-2’-deoxyguanosine (8-oxo-dG) in HCEnCs in response to O_2_ and E2. We found no significant differences in 8-oxo-dG signal in HCEnCs under any condition (*p* < 0.05, single factor ANOVA; n= 8 total, 4 females, and 4 males) (Fig 7A). Likewise, there were no significant differences when data were separated by donor sex. Interestingly, there was also no significant increase in 8-oxo-dG in HCEnCs in response to 100 μM H_2_O_2_ exposure for 48 hours. We confirmed that our experimental condition was functional by observing that 100 μM H_2_O_2_ treatment of PC3 cells for 6 hours did dramatically increase nuclear 8-oxo-dG signal (Fig 7B-C). Treatment of PC3 with H_2_O_2_ for 48 hours, like for HCEnCs, lead to almost complete death of the cells in culture.

**Fig 7.**
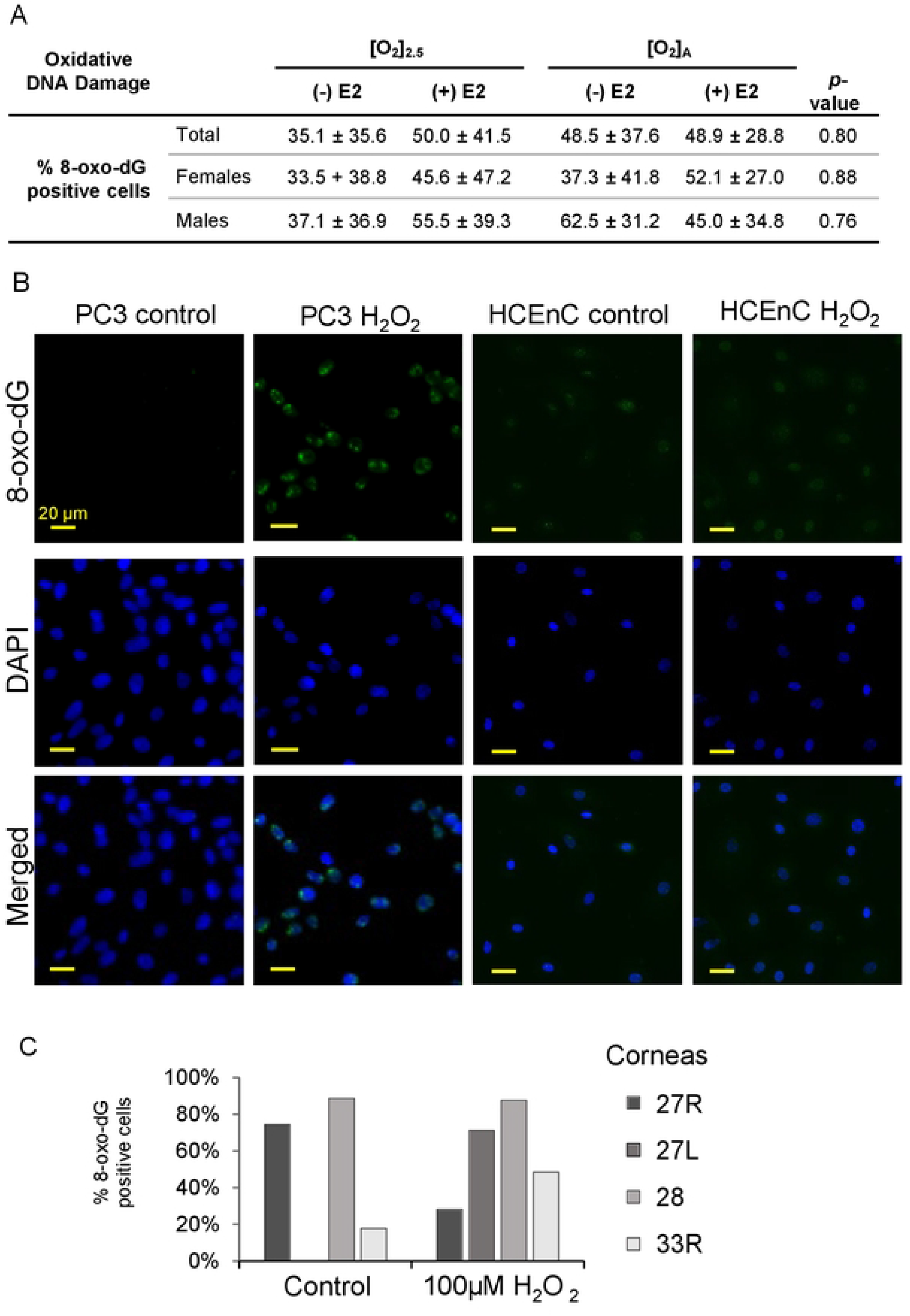
Oxidative damage in HCEnCs in the presence and absence of oxygen stress, and estradiol treatment. (A) Percentage of 8-oxo-dG positive HCEnCs in the presence or absence of estradiol treatment (10 nM) at [O_2_]_2.5_ or [O_2_]_A_. Data presented as mean ± SD. *P*-value from ANOVA single factor. N= 8 total, 4 males, and 4 females (corneas #27L-33L). (B) Immunofluorescence imaging of 8-oxo-dG in PC3 and HCEnCs following H_2_O_2_ treatment. (C) Percentage of 8-oxo-dG positive cells following H_2_O_2_ treatment in HCEnCs (n=4, corneas #27R, 27L, 28L, 33R). Donor numbers correspond to those in Table 1.

### Cellular energetics

The observed changes in mitochondrial morphology with E2 prompted us to investigate if E2 would also affect HCEnC metabolism. We measured changes in total cellular ATP levels and also evaluated for changes in oxidative respiration and glycolysis.

Cellular ATP levels were significantly different amongst the 4 conditions analyzed ([O_2_]_A_ ± E2 and [O_2_]_2.5_ ± E2; ANOVA *p*=0.01; n=7-9) (Table 3). Post-hoc analysis showed significant differences in ATP levels between cells grown at [O_2_]_A_ and [O_2_]_2.5_ without E2 (*p*=0.02). There were no significant effects of 10 nM E2 addition. However, there was 1.8-fold difference in ATP levels between cells with E2 treatment compared to cells without E2 at [O_2_]_2.5_ (*p*=0.07). When the data were segregated by sex of the donors, a cell-sex specific pattern was noted. ATP levels in female corneas showed significant differences amongst conditions (ANOVA *p*=0.01), while male corneas did not (ANOVA *p*=0.77). ATP levels in HCEnCs from female donors were higher at [O_2_]_2.5_ compared to [O_2_]_A_ (*p*=0.01) in the absence of E2.

**Table 3.** ATP levels in HCEnCs.

| | | $[O_2]_{2.5}$ | | $[O_2]_A$ | | <sup>*</sup>p-value |
| --- | --- | --- | --- | --- | --- | --- |
|  |  | (-) E2 | (+) E2 | (-) E2 | (+) E2 |  |
| **ATP (μM)** | Males | 3.85 ± 1.29 | 3.72 ± 1.38 | 3.22 ± 0.61 | 3.07 ± 1.28 | 0.77 |
| | Females | 4.83 ± 1.33 | 2.75 ± 1.64 | 2.37 ± 0.30 | 2.22 ± 0.35 | 0.009 |
| | Total | 4.46 ± 1.32 | 3.30 ± 1.45 | 2.74 ± 0.62 | 2.71 ± 1.03 | 0.01 |

Data presented as mean ± SD
<sup>\*</sup>p-value from ANOVA, n= 9 total, 4 males, and 5 females. Corneas #26R, 35R-39L.
Tukey post hoc analysis for pairwise comparisons with significant differences highlighted in red

To further characterize the effects of E2 on the energetics of HCEnC metabolism, we measured oxygen consumption rate (OCR; a measure of oxidative respiration) and extracellular acidification rate (ECAR; a measure of glycolysis) at [O_2_]_A_ and [O_2_]_2.5_ with and without E2. [O_2_]_2.5_ with and without E2 had higher OCR and ECAR measurements compared with [O_2_]_A_; however, there were no statistically significant differences in OCR or ECAR due to E2 at [O_2_]_A_ or [O_2_]_2.5_ (Fig 8A, 8B).

**Fig 8.**
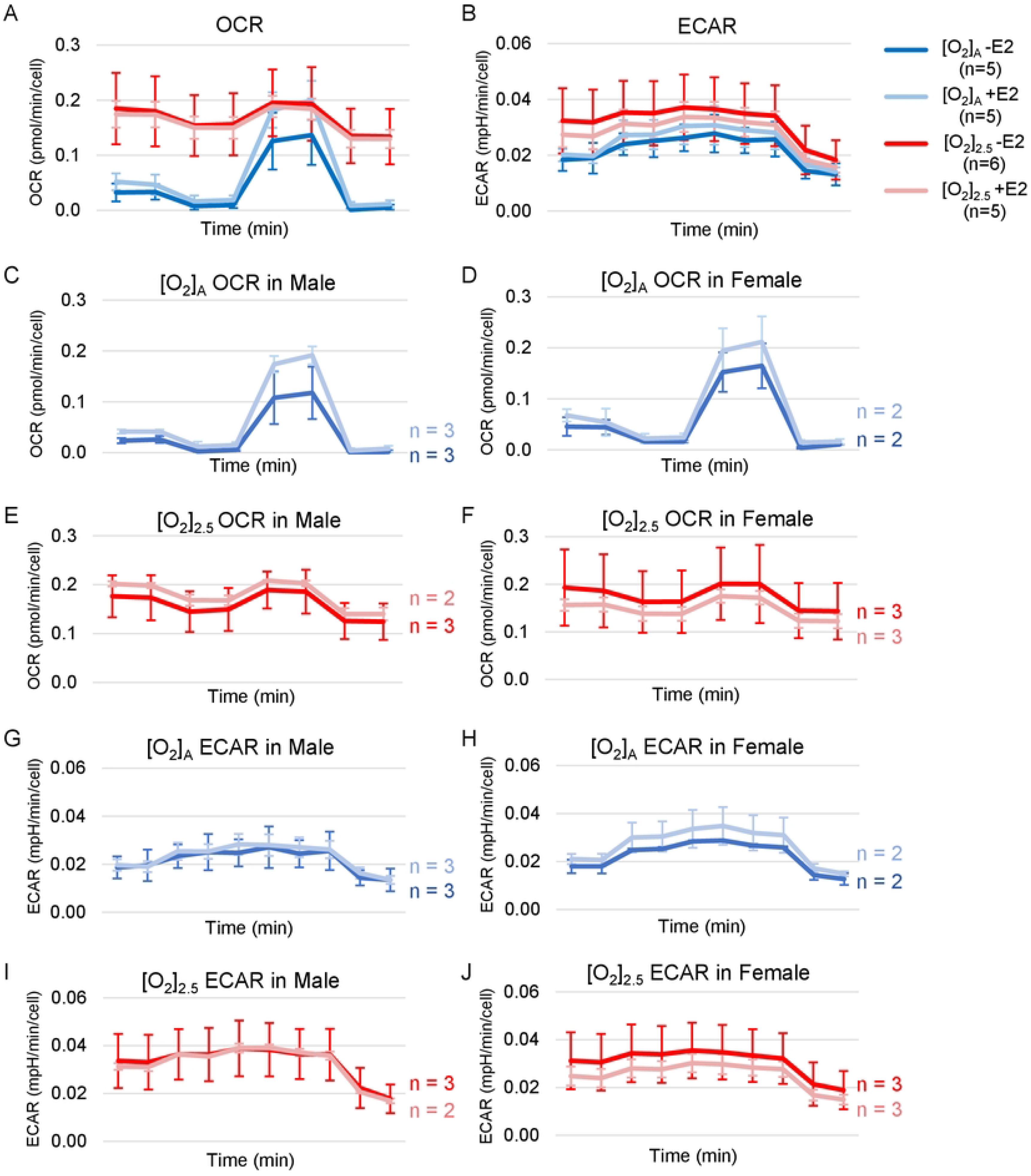
OCR and ECAR in the presence and absence of oxygen stress, and E2 treatment. (A) OCR and (B) ECAR measurements in the presence or absence of E2 treatment (10 nM) at [O_2_]_2.5_ or [O_2_]_A_. (C-J) Trends in mean data values with and without E2 treatment at both [O_2_]_2.5_ and [O_2_]_A,_ by cell sex for each condition. X-axis for time is a total duration of 41 minutes for OCR and 52 minutes for ECAR recordings. Data presented as mean ± SD. N= 5-6 total, 2-3 males, and 2-3 females (corneas #40R-45).

Because of our small sample size (n=2-3 males and 2-3 females), we did not have statistical power to identify significant differences in E2 effects by sex. However, the data trends for E2 effects by sex for each O_2_ condition are plotted in Fig 8C-J.

## DISCUSSION

The higher prevalence of FECD in women compared to men has prompted the search for a role for estrogen in FECD. Prior work has suggested toxicity of estrogen metabolites to corneal endothelium *in vitro* [35]. Yet for other chronic degenerative diseases, including ocular diseases such as macular degeneration [36, 37] and glaucoma [38-40], estrogen appears to have beneficial effects. We explored the role of estrogen and estrogen-receptor mediated effects in human corneal endothelium.

Prior data on estrogen receptor expression in corneal endothelium have been sparse and contradictory [41-43]. We found expression of ERβ and GPER, but not ERα, in HCEnC thus confirming that estrogen receptors are present in corneal endothelium. Estrogen receptors have antagonistic effects in many diseases, especially cancers, with ERα most commonly promoting cell proliferation and ERβ and GPER inhibiting tumor growth [19]. We found that E2 and G1 inhibited cell growth of the PC3 prostate cancer cell line as might be expected for cells expressing ERβ and GPER; however, we did not find statistically significant effects on HCEnCs. Since we use primary HCEnCs from older human donors, it is possible that the growth potential of our cells is limited; nevertheless, we would have anticipated seeing a robust inhibition of growth if indeed it were present.

ERβ and GPER also have significant roles in regulating cell metabolism, and estrogens mitigate the damaging effects of oxidative stress [18, 32, 44]. ERβ is the primary estrogen receptor found in mitochondria and multiple GPER-targeted interventions (pharmacologic and genetic) affect cell metabolism [45-48]. Dysregulation of cell energetics, mitochondrial dysfunction, and oxidative stress are hallmarks of FECD [31, 49, 50]. We therefore investigated measures of oxidative stress, mitochondrial dysfunction, and cell metabolism in HCEnCs under both physiologic oxygen conditions ([O_2_]_2.5_) and hyperoxic stress ([O_2_]_A_) in the presence and absence of E2. We were intrigued to find that the primary HCEnCs had stable levels of ROS and were resistant to oxidative DNA damage, even under conditions such as 100 μM H_2_O_2_ that killed PC3 cells. This appears similar to the observations of others that HCEnCs in primary cultures have the ability to restore oxidant-antioxidant balance and resist oxidative stress [51]. This is in contrast to the findings in HCEnC lines where cells are readily affected by oxidative stress [52].

Despite the absence of differences in our measures of mitochondrial morphology with hyperoxic stress in HCEnC, we detected significant effects of E2, but only for cells derived from female donors. These data support a cell-sex-specific detrimental effect of E2 when HCEnCs are exposed to hyperoxic stress. E2 and cell sex may interact in generating the sex disparity noted in FECD prevalence. It is likely that multiple stressors are required over time in order to generate the FECD phenotype [30].

However, there may also be cell-sex-dependent but E2-independent differences in HCEnC energetics, based upon our finding of different concentrations of ATP in female HCEnCs at [O_2_]_A_ vs [O_2_]_2.5_ but not in male HCEnCs. It is presently unclear whether to interpret the higher ATP levels in female HCEnCs at [O_2_]_2.5_ compared to [O_2_]_A_ as favorable or unfavorable. Based upon findings of compensatory increased mitochondrial density preceding mitochondrial burnout in HCEnCs [30], it is possible to consider that the female HCEnCs at [O_2_]_2.5_ are sick compared to male cells or the cells at the [O_2_]_A_ condition. However, overall, we have observed more favorable growth of HCEnCs at [O_2_]_2.5_ compared to [O_2_]_A_ suggesting that the cells are not sick [22]. Our data do not resolve this issue, but confirm that cell-sex does influence HCEnC behavior. Data from other cell types also support the presence of estrogen-independent mechanisms for cell-sex differences in cell behavior [33, 53].

Through these studies, we have shown that estrogen receptors ERβ and GPER are present in HCEnCs, thus providing the necessary framework for receptor-mediated estrogen signaling in HCEnCs. Furthermore, we have found that mitochondria and energetics of HCEnCs have estrogen-dependent and cell-sex-dependent effects dependent upon the O_2_ environment of the culture. We conclude that both estrogen signaling and/or cell-sex contribute to HCEnC function and dysfunction in an environment-specific fashion.

## Supporting information

**S1 Table. Primer list for quantitative RT-PCR.**

## Notes

### Competing Interest Statement

The authors have declared no competing interest.

